# Biomaterial encapsulation of human mesenchymal stromal cells modulates paracrine signaling response and enhances efficacy for treatment of established osteoarthritis

**DOI:** 10.1101/2020.07.30.228288

**Authors:** Jay M. McKinney, Krishna A. Pucha, Thanh N. Doan, Lanfang Wang, Laura D. Weinstock, Benjamin T. Tignor, Kelsey L. Fowle, Rebecca D. Levit, Levi B. Wood, Nick J. Willett

**Affiliations:** Research Division, VA Medical Center, Atlanta, GA; Department of Orthopaedics, Emory University, Atlanta, GA; Wallace H. Coulter Department of Biomedical Engineering, Georgia Institute of Technology and Emory University, Atlanta, GA; Department of Medicine, Division of Cardiology, Emory University, Atlanta, GA; Parker H. Petit Institute for Bioengineering and Bioscience, Georgia Institute of Technology, Atlanta, GA; George W. Woodruff School of Mechanical Engineering, Georgia Institute of Technology, Atlanta, GA

## Abstract

Mesenchymal stromal cells (MSCs) have shown promise as a treatment for osteoarthritis (OA); however, effective translation has been limited by numerous factors ranging from high variability and heterogeneity of hMSCs, to suboptimal delivery strategies, to poor understanding of critical quality and potency attributes. The objective of the current study was to assess the effects of biomaterial encapsulation in alginate microcapsules on human MSC (hMSC) secretion of immunomodulatory cytokines in an OA microenvironment and therapeutic efficacy in treating established OA. Lewis rats underwent Medial Meniscal Transection (MMT) surgery to induce OA. Three weeks post-surgery, after OA was established, rats received intra-articular injections of either encapsulated hMSCs or controls (saline, empty capsules, or non-encapsulated hMSCs). Six weeks post-surgery, microstructural changes in the knee joint were quantified using contrast enhanced microCT. Encapsulated hMSCs attenuated progression of OA including articular cartilage degeneration (swelling and cartilage loss) and subchondral bone remodeling (thickening and hardening). A multiplexed immunoassay panel (41 cytokines) was used to profile the *in vitro* secretome of encapsulated and non-encapsulated hMSCs in response to IL-1□, a key cytokine involved in OA. Non-encapsulated hMSCs showed an indiscriminate increase in all cytokines in response to IL-1□ while encapsulated hMSCs showed a highly targeted secretory response with increased expression of some pro-inflammatory (IL-1β, IL-6, IL-7, IL-8), anti-inflammatory (IL-1RA), and chemotactic (G-CSF, MDC, IP10) cytokines. These data show that biomaterial encapsulation using alginate microcapsules can modulate hMSC paracrine signaling in response to OA cytokines and enhance the therapeutic efficacy of the hMSCs in treating established OA.

## Introduction

Osteoarthritis (OA) is the most common chronic disease of synovial joints and is defined pathologically by degeneration of articular cartilage consisting of proteoglycan loss, chondrocyte hypertrophy, matrix fibrillation, surface erosion and lesion formation, and eventually full-thickness loss of articular cartilage resulting in bone-on-bone contact.^1,2^ OA currently impacts over 242 million people worldwide and incidence is expected to increase with rising global life expectancy.^3,4^ OA was long viewed as a degenerative disease resulting from normal body wear and tear, but the general understanding of the underlying mechanisms of OA has now expanded and it is now viewed as a multifactorial disorder that also consists of low-grade chronic inflammation.^5–7^ OA inflammation is part of a positive feedback loop that can activate chondrocytes, synoviocytes, subchondral bone cells, and other joint resident cells to secrete an array of cytokines, chemokines, and catabolic enzymes in OA.^8^ This process leads to dysregulated homeostasis of inflammatory factors including enhanced pro-inflammatory cytokine secretion [tumor necrosis factor (TNF)-α, interleukin (IL)-6, and IL-1β], which can lead to increased catabolism of the articular cartilage and increased matrix metalloproteinases (MMP) production.^9,10^ Furthermore, imbalances in anti-inflammatory cytokines [IL-1 receptor antagonist (IL-1RA), IL-4, IL-10, and IL-13] lead to attenuated chondroprotective effects and further exacerbation of OA.^11–13^ Chemokines further amplify this feedback system by stimulating neovascularization and the influx of inflammatory cells which further propagate the inflammatory response.^10,14^ Clinically, dysregulation of inflammatory cytokine balance has been shown to have a significant correlation with increased levels of knee pain.^15^ These findings have motivated the development of therapeutics that modulate the OA inflammatory disease state. Mesenchymal stromal cells (MSCs) are a promising treatment for targeting OA as they possess immunomodulatory and anti-inflammatory properties in addition to the capacity to regenerate numerous tissue types.

MSCs can be isolated from most human tissues and organs and are defined by characteristic cell surface markers and multilineage differentiation capabilities.^16,17^ While these cells do have the ability to differentiate into chondrogenic, osteogenic, and adipogenic lineages, there has been increasing interest in the paracrine signaling capabilities of MSCs as a means for their therapeutic effect in OA.^18–21^ MSC secreted factors can create a regenerative niche through numerous mechanisms, including the recruitment of additional stem and progenitor cells along with immunomodulatory effects. MSCs have the capacity to secrete an array of immunosuppressive factors [indoleamine-pyrrole 2,3-dioxygenase(IDO), TNFα-stimulated gene-6(TSG6), nitric oxide (NO), IL-10, galectins, prostaglandin E_2_ (PGE2), and transforming growth factor (TGF)-*β*] to modulate the local inflammatory environment.^22,23^ These MSC paracrine mechanisms also maintain the capacity to regulate inflammatory cell action through suppression of T-cells (proliferation and chemotaxis) and of B-cells (differentiation and chemotaxis) to further aid in modulating the inflammatory response.^23–25^ MSCs often act as sensors of the local environment and their secretome changes in response to local environmental signals; one promising approach for providing a control point for the MSC secretome is through the use of three-dimensional (3D) biomaterial constructs to deliver MSCs.

Material-based strategies have been shown to have a substantial impact on the immunomodulatory and regenerative properties of MSCs.^26,27^ The 3D environment constructed by biomaterials is well documented as a major determinant affecting MSC fate and function, and these 3D environments better replicate *in vivo* environments and cellular responses relative to two-dimensional (2D) culture systems.^28–30^ While the effects of a 3D microenvironment on MSCs has been readily studied, prior research has often focused on the effects of material-based strategies on MSC proliferation and differentiation. Encapsulation of MSCs within bulk hydrogels is a widely used strategy which can mimic native environments and allow for matrix-based signals to the cells.^30^ Biomaterial encapsulation of MSCs has been shown to increase multipotency and rates of proliferation.^27–29^ Initial work on the effects of hydrogel encapsulation on MSC immunomodulation demonstrated reduced MSC secretion of pro-inflammatory cytokines (TNF-α) and enhanced secretion of anti-inflammatory cytokines (PGE-2).^31^ Many of these same signals can be provided by cellular encapsulation in micro-sized hydrogel capsules which additionally allow for shorter diffusion distances for biochemical signals and a minimally invasive injectable means of administration.^32^ Furthermore, microencapsulation of MSCs has shown enhanced potential for suppressing proinflammatory activity of macrophages and suppressing the proliferation of peripheral blood mononuclear cells (PBMCs).^33–35^ The role biomaterials play in modulating MSC paracrine activities in the chronic inflammatory environment of OA remains understudied, especially *in vivo*.

In the current study, sodium alginate cellular encapsulation was used to modulate the paracrine signaling properties of human MSCs (hMSCs). Alginate is a heteropolysaccharide which is used in many biomedical applications; it is highly tailorable and can be modified to yield variable levels of biodegradability, mechanical strength, and cellular affinity, among numerous other properties.^36^ In previous work we utilized an alginate microencapsulation system with 1% ultrapure low viscosity sodium alginate crosslinked with BaCl_2_ to create microcapsules approximately 150 μm in diameter.^18,37,38^ These studies demonstrated that this encapsulation system permits molecules < 80 kDa to diffuse in and out while preventing the MSCs from integrating with the host tissue.^37^ Furthermore, in an OA rat model we showed that encapsulated hMSC viability following intra-articular injection was ~ 9 days while nonencapsulated hMSC remained viable for ~ 7 days.^18^ We also demonstrated that intra-articular injection of encapsulated hMSCs ameliorated the onset of post traumatic OA.^18^ To further explore the utility of this encapsulation system, it was used in the current study to analyze the therapeutic efficacy of encapsulated hMSC in established OA.

While pre-clinical OA studies have shown MSC treatment can be chondroprotective (attenuating cartilage breakdown), MSCs have yet to translate into a consistently reliable and effective clinical therapy.^39^ A distinct gap in the pre-clinical literature is that the majority of the work studying MSC efficacy in OA focuses on preventing disease development; however, in clinical scenarios, patients that are treated with cell therapies typically have already developed OA when they seek treatment.^22^ There is a fundamental difference in the local environment, and likely the necessary therapeutic mechanism of action, between an injured joint prior to OA development (but primed to develop OA) and a joint where OA is established. Thus, there is a substantial gap in our understanding of the therapeutic efficacy of MSCs in delaying further disease progression once OA is established. The objective of the current study was to assess the effects of biomaterial encapsulation in alginate microcapsules on hMSC secretion of immunomodulatory cytokines in an OA microenvironment and therapeutic efficacy in treating established OA.

## Materials and methods

### Cell culture

Bone marrow derived hMSCs, obtained from the Emory Personalized Cell Therapy Core (EPIC) facility at Emory University, were cultured in complete minimum essential medium Eagle-α modification (α-MEM; 12561; Gibco, Carlsbad, CA, USA) supplemented with 10% heat-inactivated fetal bovine serum (FBS; S11110H; Atlanta Biologicals, Lawrenceville, GA, USA), 2 mM L-glutamine (SH3003401; HyClone, Logan, UT, USA), and 100 μg/mL penicillin/streptomycin (P/S; B21110; Atlanta Biologicals, Lawrenceville, GA, USA), and subcultured at 80% confluency. The three criteria for hMSC designation – tri-lineage differentiation, surface marker phenotyping, and adherence to plastic – were confirmed in a previous study.^18^ Briefly, differentiation was confirmed for chondrogenesis, adipogenesis, and osteogenesis; hMSC phenotyping confirmed cells were positive for MSC markers, including CD73, CD90, and CD105, and negative for hematopoietic markers, including CD45, CD34, CD11b, CD79A, and HLA-DR; expansion of hMSCs relied on adherence to plastic tissue culture plates up to passage 4.

### Cell encapsulation

An electrostatic encapsulator was used to encapsulate 5 × 10^5^ cells/mL hMSCs (passage 4) suspended in 2% ultrapure low viscosity sodium alginate LVG (UP-LVG; 4200006; PRONOVA™ UP LVG; NovaMatrix, Sandvika, Norway) using the following parameters: 0.2 μm nozzle, 2.5 mL/h flow rate, and 7 kV voltage. Capsules were gelled in 50 mM BaCl_2_ and subsequently washed two times with 0.9% saline (NaCl), and re-suspended to the appropriate dose in complete α-MEM (*in vitro* hMSC culture model) or saline (*in vivo* MMT model). A Live/ Dead™ Viability/Cytotoxicity kit (L3224; Invitrogen, Carlsbad, CA, USA) was used to assess encapsulated hMSC viability. The average diameter of encapsulated hMSC microspheres was 144 ± 16 μm, as previously quantified.^18^ Empty capsules were manufactured using the same procedure, without the addition of hMSCs, and yielded similar size.

### In vivo MMT model

All animal care and experiments were conducted in accordance with the institutional guidelines and approved by the Atlanta Veteran Affairs Medical Center (VAMC) with experimental procedures approved by the Atlanta VAMC Institutional Animal Care and Use Committee (IACUC). Weight-matched (300-350 g) male Lewis rats (strain code: 004; Charles River, Wilmington, MA, USA) were used for the medial meniscal transection (MMT) model used to induce OA in the current study.^40^ Briefly, animals were anesthetized under isoflurane and injected subcutaneously with 1 mg/kg sustained-release (SR) buprenorphine (ZooPharm, Windsor, CO, USA). The skin over the medial aspect of the left femoro-tibial joint was sterilized, and a blunt dissection was used to expose the medial collateral ligament (MCL) and transect it to expose the meniscus. A full-thickness cut was made through the meniscus at its narrowest point followed by soft tissue re-approximation and closure using 4.0 Vicryl sutures and wound clips for skin closure. Sham surgery involved MCL transection, with no transection of the meniscus, followed by closure of the skin. Intra-articular injections were performed at three weeks post-surgery, the time point in the MMT model corresponding to the presentation of OA-associated cartilage degeneration and osteophyte formation.^40,41^ Intra-articular injections were performed using a 25-gauge needle and included: hanks balanced salt solution (HBSS; MMT/Saline; n = 7), empty sodium alginate capsules in HBSS (MMT/Empty Caps; n = 7), 5×10^5^ non-encapsulated hMSC in HBSS (MMT/hMSC; n = 8) and 5×10^5^ encapsulated hMSC in HBSS (MMT/Encap hMSC; n=7). Rats in the MMT/Encap hMSC were injected within two hours of hMSC encapsulation (cells were stored at 4°C until injection). Sham (n = 6) animals did not receive any injection post-surgery. At six weeks post-surgery, animals were euthanized by CO_2_ inhalation. Left hind limbs were collected and fixed in 10% neutral buffered formalin for a two-day minimum before further sample preparation.

### microCT analysis of articular joint parameters

Prior to scanning, all muscle and connective tissue from collected hind limbs was removed, the femur was disarticulated from the tibia, and all peripheral connective tissue surrounding the joint was removed to expose the articular cartilage of the medial tibial plateau. Tibiae were immersed in 30% (diluted in PBS) hexabrix 320 contrast reagent (NDC 67684-5505-5, Guerbet, Villepinte, France) at 37°C for 30 minutes before being scanned.^42^ All samples were scanned using equilibrium partitioning of an ionic contrast agent based micro-computed tomography (EPIC-μCT; microCT) through the use of a Scanco μCT 40 (Scanco Medical, Brüttisellen, Switzerland) using the following parameters: 45 kVp, 177 μA, 200 ms integration time, isotropic 16 μm voxel size, and ~27 min scan time.^42^ Scans were read out as 2D tomograms which were subsequently orthogonally transposed to yield 3D reconstructions for all scanned samples. All microCT parameters (articular cartilage, osteophyte, and subchondral bone) were evaluated as previously described.^43,44^ For cartilage parameters, thresholding of 110 - 435 mg hydroxyapatite per cubic centimeter (mg HA/cm^3^) was used to isolate the cartilage from the surrounding air and bone. Furthermore, for bone parameters, thresholds of 435 - 1200 mg HA/cm^3^ were implemented to isolate bone from the overlying cartilage. Coronal sections were both evaluated along the full length of the cartilage surface (total) and in third (medial, central, and lateral) regions of the medial tibial condyle. For articular cartilage, volume, thickness, and attenuation parameters were quantified. Attenuation is inversely related to sulfated glycosaminoglycans (sGAG) content.^42^ In OA, sGAG concentration in articular cartilage decreases due to degradation, creating a gradient which leads to an increased hexabrix concentration in the cartilage. High hexabrix and low sGAG levels (increased sGAG loss) correspond to a higher attenuation value. In addition to microCT analysis of articular cartilage, osteophyte volumes found on the most medial aspect of the medial tibial plateau were evaluated for their cartilaginous and mineralized portions. Additionally, subchondral bone was evaluated for volume, thickness, and attenuation (indirect measure of bone mineral density) along the total, medial, central, and lateral regions, similar to the approach used for articular cartilage analysis.

### Surface roughness of articular cartilage

Articular cartilage surface roughness and exposed bone surface area (full thickness lesion surface area) were quantified using a custom MATLAB script, *SurfaceRoughness* function.^45^ Briefly, coronal sections from the Scanco μCT 40 were exported as 2D TIFF images and imported into MATLAB R2016a (MathWorks, Natick, MA, USA). A custom code created a 3D surface with these images by scanning section images sequentially and consolidating them. This 3D surface was fitted along a computationally generated 3D polynomial surface, unique for each sample imported, which was fourth order along the ventral-dorsal axis and second order along the medial-lateral axis. Surface roughness was quantified as the root mean square of differences between the 3D surface created with the exported TIFF images and the polynomial fitted surface. Exposed bone (full thickness lesion surface area) was quantified as root mean square of area where the difference between the 3D cartilage surface and 3D bone surface was ≤ three pixels (i.e. no presence of any articular cartilage). Threshold values of 110 - 435 mg HA/cm^3^ were used to separate cartilage from air and the underlying subchondral bone and threshold values of 435 - 1200 mg HA/cm^3^ were used to isolate bone from overlying cartilage (matching microCT thresholds). All MATLAB analyses were performed along the total surface and in regions (medial, central, lateral) of the medial tibial plateau, similar to the microCT analyses.

### Histology

To prepare bone samples for sectioning, tibiae were decalcified in Immunocal (SKU-1414-32; StatLab, McKinney, TX, USA) for 14-21 days. Dehydrated samples were processed into paraffin-embedded blocks, sectioned at 5 μm thickness and stained. Hematoxylin and eosin (H&E; Fisherbrand™ 517-28-2, Waltham, MA, USA), and Safranin O and fast green (Saf-O; Electron Microscopy Sciences^®^ 20800, Hatfield, PA, USA) were used to stain all study samples in accordance with manufacturer protocols. For all samples, a single representative image was selected for H&E and Saf-O (serial sections).

### In vitro hMSC cytokine analysis model

Passage 4 hMSCs, matching donor with *in vivo* MMT model, were utilized *in vitro*. Nonencapsulated hMSCs were sub-cultured to 80% confluency in complete α-MEM medium in 12-well plates and cultured at 37°C, 5% CO_2_. For encapsulated hMSCs, immediately following encapsulation and washing, cells were placed in treatment medium (unstimulated or stimulated) in 12-well plates at 37°C, 5% CO_2_. Unstimulated media (+CTRL) contained complete α-MEM medium only and stimulated media (+IL-1β) contained 20 ng/mL IL-1β (FHC05510, Promega, Madison, WI, USA) in complete α-MEM medium. IL-1β was used to model the OA inflammatory environment in the current study as it is a major pro-inflammatory modulator in OA.^9^ IL-1β concentration (and group sample size) were selected based on prior experiments and preliminary data.^46,47^ Stimulated and unstimulated media were added to non-encapsulated and encapsulated hMSCs at day 0 (n=6) followed by a 24-hour stimulation period with media collection at the end point for stimulation for the four study groups. Additional filtering steps were implemented in order to remove encapsulated hMSCs by passing collected media through a 9 μm filter. Samples were stored at −80°C until Luminex analysis was performed. Loaded samples (2.03 μL) were determined to be within the linear range of detection of the MAGPIX (MAGPIX-XPON4.1-CEIVD; EMD Millipore Corporation, Burlington, MA, USA) system. Cytokines were quantified using a bead based multiplex immunoassay, Luminex Cytokine/Chemokine 41 Plex Kit (HCYTMAG-60K-PX41; EMD Millipore Corporation, Burlington, MA, USA). Median fluorescent intensity values were read out using Luminex xPONENT software V4.3 in the MAGPIX system. Background subtraction was performed on non-stimulated and stimulated conditions using read out values from media only and 20 ng/mL IL-1β supplemented media, respectively.

### Partial Least Squares Regression (PLSR)

Partial least squares discriminant analysis (PLSDA) was performed in MATLAB (Mathworks, Natick, MA, USA) using a function written by Cleiton Nunes (Mathworks File Exchange).^48^ This approach accounts for the multivariate nature of the data without overfitting.^49,50^ Prior to inputting the data into the algorithm, all cytokines were z-scored [(observed - mean) / standard deviation]. For articular cartilage, osteophyte, and subchondral bone *in vivo* analyses, total and medial microCT parameters were used as the independent variables and the five separate treatment groups were used as the outcome variables. For cytokine analysis of the *in vitro* cell culture model, cytokine measurements were used as the independent variables and the four individual treatment groups were used as the outcome variable. Latent variables (LV) in a multidimensional space (dimensionality varied by number of independent input variables) were defined and the two primary LVs were used for orthogonal rotation to best separate groups in the new plane defined by LV1 and LV2 (Fig. 1a). In the *in vitro* cell culture model orthogonality between encapsulated hMSCs (LV1 horizontal axis) and non-encapsulated hMSCs (LV2 vertical access) was confirmed via dot product (LV1 o LV2 = - 1.059 x 10-16 ~ 0). Loadings plots were generated from this analysis and display the relative importance of input variables (microCT parameters or cytokines) in contributing to the final composite values (scores) for each sample (Fig. 1b&c). Error bars on each cytokine (in the loadings plots) were computed by PLSR model regeneration using iterative (1000 iterations) leave one out cross validation (LOOCV). To further confirm significant differences between groups assessed in PLSDA, the true differences in centroids (center of mass) of all groups were compared against the differences computed by a random distribution obtained by permuting the group labels 100 times. For each test, true group assignment showed p_permute_<0.05 compared to the randomly permuted distribution, further confirming the validity of the data.

**Fig. 1.**
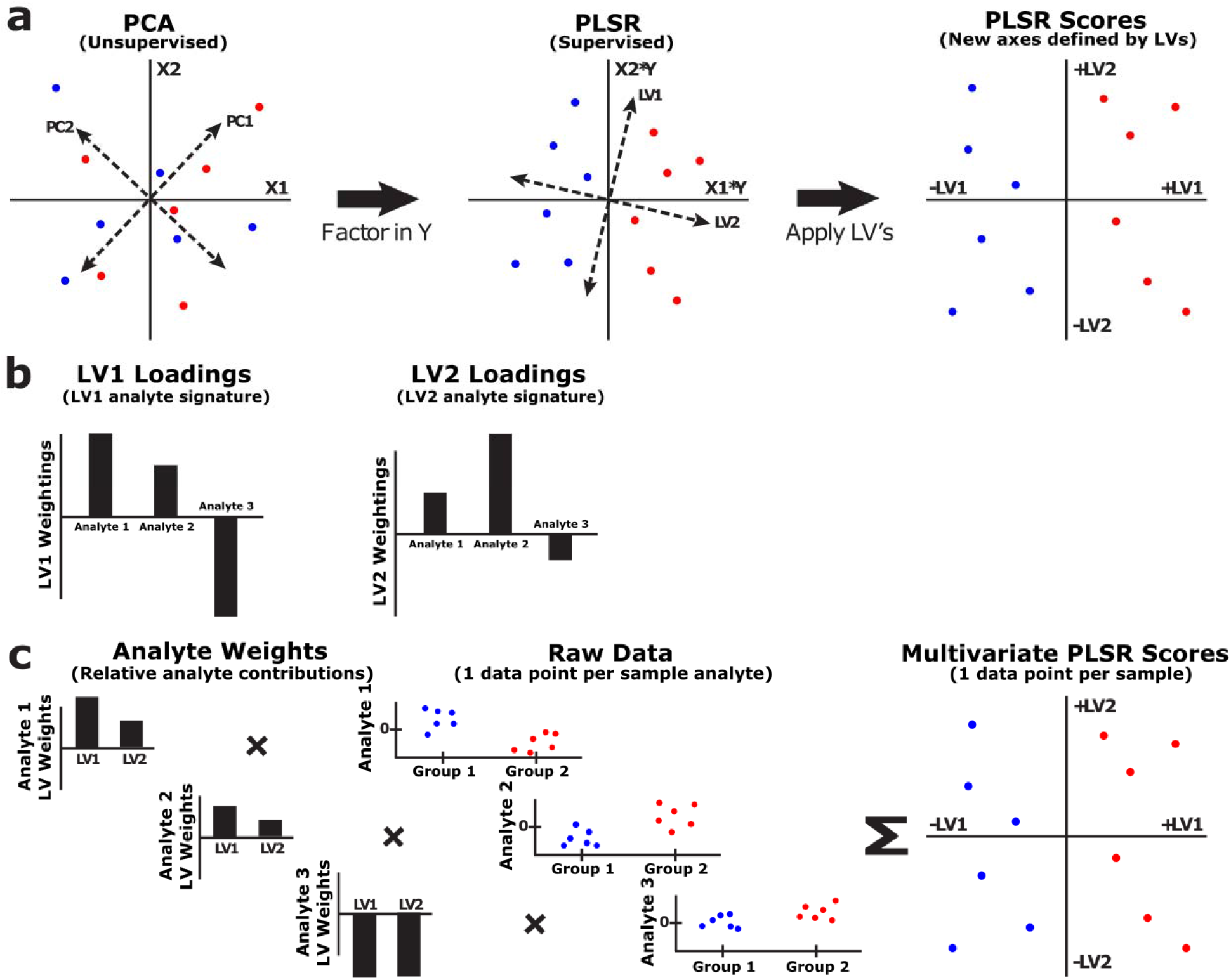
(a) Principle component analysis (PCA) identifies axes of maximum variation among samples in the data when measurement variables (**X1** and **X2**) are plotted against one another. Through incorporation of a response variable **Y**, partial least squares regression (PLSR), enables identification of maximum co-variation between the **X** variables and different **Y** responses. PLSR outputs new linear combinations of **X** variables, referred to as latent variable**s** (LVs). (b) Each latent variable is comprised of weights, which ranks the importance of each input variable X_i_, in determining the final composite values for each sample data point. (c) To obtain the PLSR scores plot, the raw data is multiplied by the calculated weights for each latent variable (**LV1** and **LV2**). The new axes defined by these latent variables (**LV1** and **LV2**) better separates the data with respect to the identity of the **Y** response variables.

### Statistics

All data is presented as mean ± standard deviation (SD). Significance for all microCT parameters was determined with one-way ANOVA with post hoc Tukey honest test for articular cartilage and subchondral bone parameters. Bonferroni correction was used for post hoc analysis for the exposed bone and osteophyte parameters due to their nonparametric nature. For all PLSDA scores plots, significance was determined with one-way ANOVA and post hoc Tukey honest test. To determine significant differences between encapsulated hMSCs in unstimulated (+CTRL) and stimulated (+IL-1β) conditions, two tailed t-tests were used with

Bonferroni correction to account for the independent analysis of multiple groups. Statistical significance was set at p < 0.05. All data were analyzed using the *R stats, ggsignif*, and *ggpubr* packages in R (The R Foundation, Vienna, Austria).

## Results and discussion

### Qualitative analysis of the therapeutic efficacy of encapsulated hMSCs

While MSCs have been readily studied in the context of OA, the major focus of previous research has targeted delaying OA development.^22,51^ To address this gap in knowledge, the effect of MSCs in delaying further disease progression of established OA was assessed in the current study. This approach provides added clinical relevance since clinical OA is more commonly diagnosed based on progressive disease phenotypes, including joint space narrowing and osteophyte development, as pain is commonly associated with these more established manifestations of the disease leading patients to seek medical treatment.^52,53^ The rat MMT OA model was used as an OA phenotype manifests by the 3 week timepoint and further progression is observed at the six week timepoint; encapsulated hMSC treatment was administered at three weeks post-surgery, once OA had already been established, and the effects of the treatment were evaluated at the six week endpoint.^44^ At the time of injection, encapsulated hMSC viability was 96.4 ± 2.1%. Encapsulation was leveraged in the current study to assess its effects on modulating the paracrine response of hMSCs in established OA.

Histology (H&E and Saf-O) was performed on tibiae to qualitatively assess the effects of encapsulated hMSC therapeutics on OA progression (Fig. 2a-j). Representative histological images of the Sham group showed consistent proteoglycan staining and a smoothness along the entire medial articular cartilaginous surface and no presence of osteophyte development on the most medial aspect of the joint (Fig. 2a&f). All MMT conditions showed proteoglycan loss (loss of Saf-O staining), loss of chondrocytes (lack of hematoxylin staining in certain regions of the articular cartilage layer), presence of fibrillations in the articular cartilage layer, and the development of osteophytes (Fig. 2b-e&g-j). While all MMT conditions did show variable levels of cartilage damage, the MMT/Encap group showed qualitatively less cartilage degeneration and surface roughness, relative to all other MMT conditions (Fig. 2e&j). Furthermore, while all MMT conditions showed osteophytes developing on the marginal edges of the joint, the MMT/hMSC and MMT/Encap hMSC group showed qualitatively larger areas relative to the MMT/Saline and MMT/Empty Caps group (Fig. 2d-e&i-j). Representative cartilage surface renderings were generated for each sample from each group using microCT captured images (samples matched with representative histology; Fig. 2k-o). Analysis of cartilage surface roughness was performed by subtracting individual 3D polynomial surfaces from the corresponding cartilage surface renderings (generated by custom MATLAB algorithm). All MMT conditions exhibited changes in articular cartilage structure as can be visualized with the development of elevations (red regions) and depressions (blue regions) relative to the Sham group. Furthermore, the MMT/Encap hMSC showed qualitatively attenuated cartilage surface roughness relative to other MMT groups (Fig. 2k-o). Together, these metrics demonstrated that with encapsulated hMSC treatment there was qualitatively less cartilage degeneration when compared to the other MMT groups.

**Fig. 2.**
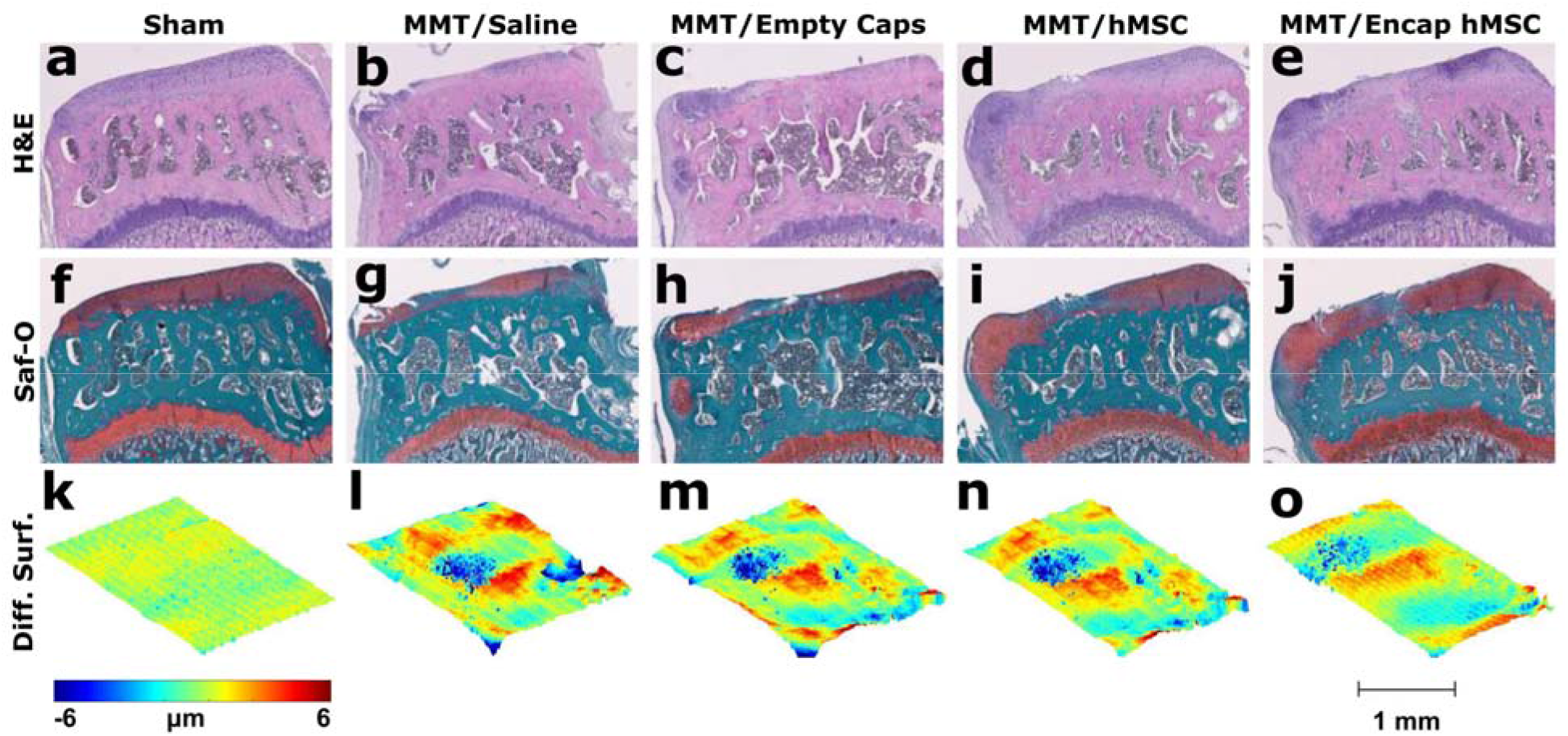
(a-j) Serial hematoxylin and eosin (H&E; a-e) and safranin-O and fast green (Saf-O; f-j) coronal sections of rat medial tibial plateaus at six weeks after Sham or MMT operation of rat hindlimbs. For MMT-induced OA there is presence of increased articular cartilage degeneratio**n** [increased proteoglycan loss (loss of red coloration in Saf-O images), loss of articular chondrocytes (lack of hematoxylin stain), surface fibrillations, formation of erosions and lesion**s]** and the presence of osteophyte formations on the marginal edges for all MMT conditions (b-e&g-j). Sham operated hindlimbs (a&f) did not show any damage to articular cartilage or the presence of osteophyte formations. The MMT/Encap hMSC group (e&j) showed less overall cartilage damage (smoother cartilage surface with less erosion and lesion formation) with respect to all other MMT conditions. (k-o) MATLAB generated representative topographic maps of the articular cartilage surfaces depict the deviation of a sample’s cartilage surface from a 3D polynomial fitted surface. Representative surface renderings were matched with representative histology and microCT. These surface renderings demonstrate elevations (red) and depressio**ns** (blue) in articular cartilage surfaces of MMT groups which were not found in the Sham. All images are oriented with the medial aspect of the tibia on the left. Scale bar (bottom right corner) is universal for all histology representative images. Scale bar for the topographic surfa**ce** renderings (bottom center) is also included.

### Encapsulated hMSCs attenuated cartilage degeneration in established OA

For quantitative analysis, microCT was employed to study the articular cartilage, osteophytes, and subchondral bone regions of rat tibiae, as previously described.^43,44^ MicroCT analysis of OA phenotypes has been demonstrated to be comparable to histopathology – the gold standard reference for OA characterization – when assessed in 2D and more sensitive than histopathology when used to assess 3D parameters.^43^

Detailed analysis of the articular cartilage was performed on various morphological parameters for both the total and segmented regions (medial, central, and lateral) of the medial tibial condyle (Fig. 3&S1). Previous studies have demonstrated that OA development in the MMT model is largely localized to the medial plateau and specifically the medial 1/3 region of the articular cartilage.^44^ For total cartilage volume, all MMT conditions showed elevated cartilage volume relative to Sham (Fig. 3a). Treatment with MMT/Encap hMSCs attenuated the MMT induced increase in cartilage volume that was found in the MMT/Saline and MMT/Empty Caps groups. However, no significant difference was found between MMT/Encap hMSC and MMT/hMSC alone, suggesting a milder effect of non-encapsulated cells relative to the encapsulated hMSCs. For medial cartilage volume analysis, significant increases in volume were yielded for the MMT/Saline, MMT/Empty Caps, and MMT hMSC groups. The MMT/Encap hMSC group did not show significant increases in cartilage volume relative to Sham (Fig. 3b). Cartilage thickness yielded similar outcomes to the volume parameter as no significant difference for cartilage thickness were found between the MMT/Encap hMSC group relative to the Sham group for both total and medial analysis (Fig. 3c&d). Furthermore, for the medial thickness parameter there were no significant differences noted between MMT/Encap hMSC and MMT/hMSC, further demonstrating the mild therapeutic effect of the non-encapsulated hMSCs. Cartilage attenuation, which permits the indirect quantification of proteoglycan content, yielded a single significant difference between Sham and MMT/hMSC for total analysis (Fig. 3e). While the medial analysis did discern differences between the Sham and all MMT conditions, there were no differences found between the MMT conditions as they all demonstrated increased attenuation (decreased proteoglycan content) relative to the Sham group (Fig. 3f). Additional analysis of the articular cartilage surface was performed using a custom MATLAB script to quantify surface roughness and exposed bone surface area (full thickness lesion surface area).

**Fig. 3.**
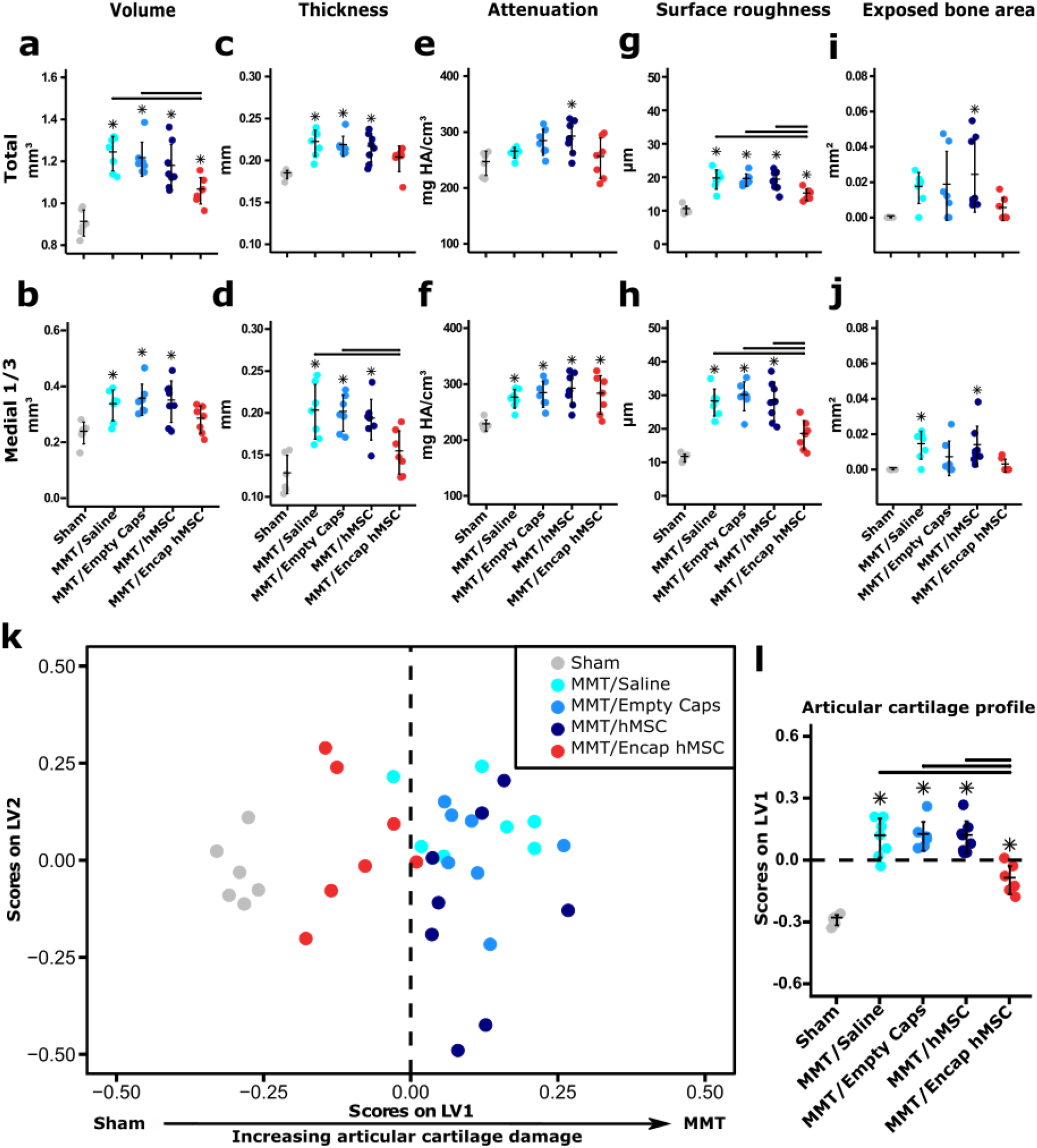
(a) Total articular cartilage volume for all MMT groups are significantly greater than Sham animals; total articular cartilage volume for MMT/Encap hMSC group is significantly lower than MMT/Saline and MMT/Empty Cap groups. (b) Medial 1/3 articular cartilage volumes for MMT/Saline, MMT/Empty Caps and MMT/hMSC groups were significantly greater than Sham; hMSC/Encap hMSC group was not significantly different from Sham. (c&d) Total and medial 1/3 articular cartilage thickness values for MMT/Saline, MMT/Empty Caps and MMT/hMSC groups were significantly greater than Sham; no differences were found for either parameter between MMT/Encap hMSC and Sham; medial 1/3 articular cartilage thickness for MMT/Encap hMSC group was significantly lower than MMT/Saline and MMT/Empty Caps groups. (e) Total articular cartilage attenuation for the MMT/hMSC group was significantly greater than the Sham group. (f) Medial 1/3 cartilage attenuation values for all MMT groups were significantly greater than the Sham group and no differences were found among MMT conditions. (g) Total cartilage surface roughness showed significantly higher values for all MMT groups relative to Sham; the MMT/Encap hMSC group did show significantly less surface roughness than all other MMT conditions. (h) Medial 1/3 analysis of surface roughness yielded identical findings to total analysis except no difference was found between Sham and MMT/Encap hMSC groups. (i&j) A difference in total exposed bone was found only for the MMT/hMSC group compared to all other groups; for medial 1/3 analysis of exposed bone, MMT/Saline and MMT/hMSC groups were significantly increased from Sham group. (k) PLSDA assessment of the overall effect of the therapeutics applied on articular cartilage damage showed distinct separation with Sham and MMT Encap hMSC separating to the left and all other MMT conditions separating to the right along LV1. (l) Quantification of the scores obtained from PLSDA analysis demonstrated that all MMT conditions had significantly more cartilage damage than Sham; in addition, MMT/Encap hMSC had significantly less damage than all other MMT conditions. Data presented as mean +/- SD. *n =* 6 for Sham, *n* = 7 for MMT/Saline, *n* = 7 for MMT/Empty Caps, *n* = 8 for MMT/hMSC and *n* = 7 for MMT/Encap hMSC. * represents significant differences (p < 0.05) between individual MMT conditions and Sham. Horizontal black bars indicate significance (p < 0.05) between individual MMT groups.

Surface roughness analysis of the articular cartilage surface provides a quantitative measure of changes that may arise from matrix fibrillation, erosion and lesion formation, and full-thickness cartilage loss (exposed bone surface area). All MMT conditions showed significantly increased surface roughness relative to Sham for analysis of the total tibial plateau (Fig. 3g). For medial surface roughness analysis, the MMT/Saline, MMT/Empty Caps, and MMT/hMSC groups showed significantly increased surface roughness compared to Sham but no difference was detected between MMT/Encap hMSC and Sham (Fig. 3h). Additionally, treatment with MMT/Encap attenuated the surface roughness to a level significantly lower than the other MMT conditions for both total and medial analyses (Fig. 3g&h). To further characterize changes to articular cartilage, exposed bone surface area was quantified. The total surface area of exposed bone was significantly different between the MMT/hMSC group and the Sham group (Fig. 3i). For medial analysis of exposed bone, both MMT/Saline and MMT/hMSC showed a significant increase relative to Sham (Fig. 3j). Additionally, incidence of exposed bone was assessed by group: MMT/Encap hMSC had the least number of samples with exposed bone (4/7), followed by MMT/Empty Caps (5/7), MMT/Saline (6/7), and MMT/hMSC (8/8). Fisher’s exact test was performed for contingency analysis of exposed bone groups. MMT/Encap hMSC showed no significant difference from Sham, whereas all other MMT groups were significantly different from Sham (0/6; Fig. S2a). For the analysis of incidence of exposed bone on the medial aspect of the joint, increased exposed bone incidence was found in both MMT/Saline (5/7) and MMT/hMSC (8/8) groups relative to Sham (0/6; Fig. S2b). No significant differences were found between Sham and MMT/Encap hMSC (3/7) or MMT/Empty Caps (4/7) groups (Fig. S2b).

To assess the overall effect of encapsulated hMSCs on MMT induction, factoring in all articular cartilage parameters (total and medial) as model inputs, PLSDA was implemented to identify new axes which better separate the data with respect to the identity of the treatment group (Sham and MMT groups). LV1 separated out groups by severity of cartilage damage with the Sham and MMT/Encap hMSC groups separating to the left from the MMT/Saline, MMT/Empty Caps, and MMT/hMSC groups, on the right (Fig. 3k). A one-way ANOVA of the

LV1 scores demonstrated that all MMT conditions had significantly higher scores (increased damage) than the Sham; while the MMT/Encap hMSC group was significantly lower than the other MMT groups (Fig. 3l). However, MMT/Encap hMSC also demonstrated a higher LV1 score compared to the Sham, suggesting increased cartilage damage (Fig. 3l). Overall, these metrics reveal that encapsulated hMSCs provided a positive therapeutic protective effect on articular cartilage in established OA.

Articular cartilage degeneration, the primary outcome of OA, manifests with proteoglycan loss, resulting in increased water concentration in cartilage (swelling), superficial cartilage matrix fibrillation, erosion and lesion formation, and eventually full-thickness loss of articular cartilage resulting in bone-on-bone contact.^2,54^ Further development of these articular cartilage phenotypes was attenuated with encapsulated hMSC treatment and not found with hMSCs or alginate (empty capsules) alone. Specifically, encapsulated hMSCs attenuated further articular cartilage swelling (volume and thickness) and delayed the further development of matrix fibrillations and erosion and lesion formation (surface roughness). Additionally, there was lower incidence of full thickness cartilage loss (exposed bone) after treatment with encapsulated hMSCs. These findings suggest that while encapsulated hMSCs elicit a positive therapeutic effect in delaying further articular cartilage degeneration, this treatment did not fully restore articular cartilage to a similar state as the Sham control (regenerative effect). There was still significant cartilage proteoglycan loss (attenuation), cartilage swelling, and full thickness cartilage loss; these levels were comparable to previous reports of the disease state at the three week time point (i.e. the time point of the encapsulated hMSC intervention).^18,43–45,55^ While encapsulated hMSCs may not necessarily have any restorative properties that can reverse established disease damage, this treatment does have disease modifying potential that still provides high clinical translatability.

### Encapsulated hMSCs had a minimal therapeutic effect on subchondral bone remodeling in established OA

Subchondral bone, which is the bone layer immediately underlying articular cartilage, has been observed to harden and thicken during the progression of OA (sclerosis), especially at later stages of the disease.^53,56–58^ Subchondral bone sclerosis, along with joint space narrowing, is one of the most common phenotypes observed in clinical OA via standard x-ray diagnostic imaging.^52,53,57^

Detailed analysis of subchondral bone was performed on various morphological parameters for both the total and segmented regions (medial, central, and lateral) of the medial tibial condyle (Fig. 4&S3). Previous work with the rat MMT model at three weeks post-surgery has shown that there is increased incidence of subchondral bone remodeling in the medial 1/3 region.^59^ Subchondral bone volume, for both total and medial regions, showed MMT/Saline, MMT/Empty Caps, and MMT/hMSC groups were significantly elevated relative to Sham (Fig. 4a&b). For the subchondral bone thickness parameter, all MMT conditions demonstrated significantly increased values relative to Sham (Fig. 4c&d). Total attenuation was not significantly different between MMT/Encap hMSC and Sham or between MMT/hMSC and Sham (Fig. 4e). The other two MMT conditions (MMT/Saline and MMT/Empty Caps) had a significant increase in attenuation values relative to the Sham group, indicating increased bone mineral density (hardening) of the subchondral bone (Fig. 4e). However, the analysis of attenuation for the medial 1/3 plateau showed significantly higher values for all MMT groups compared to Sham (Fig. 4f).

**Fig. 4.**
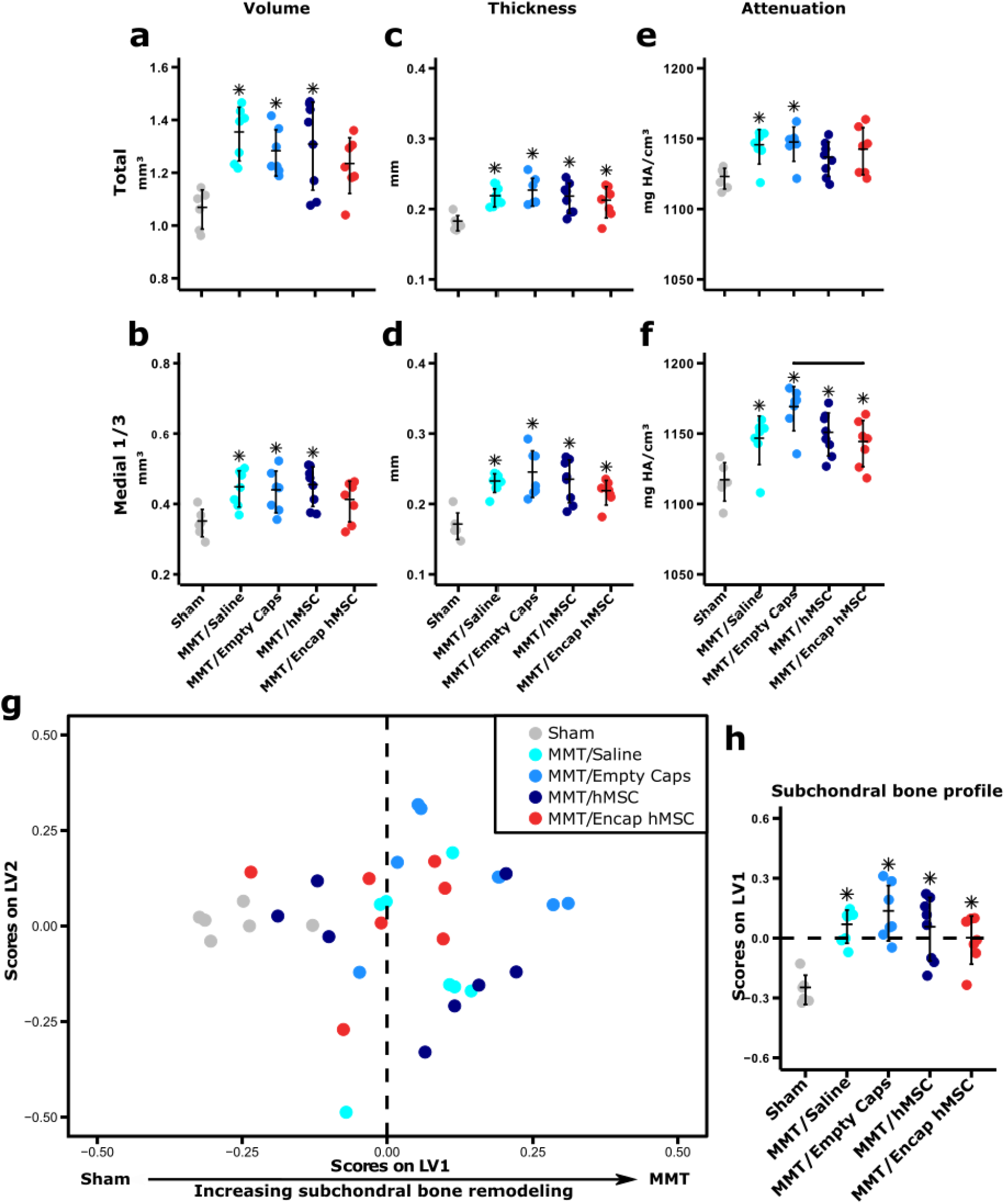
(a&b) Total and medial 1/3 subchondral bone volumes for all MMT groups, except MMT/Encap hMSC, were significantly greater than the Sham group. (c&d) Total and medial 1/3 subchondral bone thickness analysis yielded significant increases in all MMT groups relative to Sham. (e) MMT/Empty Caps and MMT/Saline total subchondral bone attenuation was significantly greater than the Sham while showing no differences with MMT/hMSC and MMT/Encap hMSC groups. (f) In the medial 1/3 region, all MMT groups had significantly greater attenuation values compared to Shams; the only difference found between MMT conditions was between MMT/Empty Caps and MMT/Encap hMSC. (g) PLSDA analysis of total and medial 1/3 subchondral bone parameters depicted significant separation between Sham, to the left, from all MMT conditions, to the right, along LV1 based on the level of subchondral bone remodeling. (h) Statistical analysis of these scores demonstrated a significant difference between the Sham group and all the MMT groups, with no differences between the respective MMT groups. Data presented as mean +/- SD. *n =* 6 for Sham, *n* = 7 for MMT/Saline, *n* = 7 for MMT/Empty Caps, *n* = 8 for MMT/hMSC and *n* = 7 for MMT/Encap hMSC. * represents significant differences (p < 0.05) between individual MMT conditions and Sham. Horizontal black bars indicate significance (p < 0.05) between individual MMT groups.

Cumulative analysis of the efficacy of encapsulated hMSCs on the subchondral bone layer in OA (accounting for volume, thickness, and attenuation parameters) was analyzed with PLSDA. LV1 separated out all study groups by levels of subchondral bone remodeling with Sham separating out on the left from all MMT groups on the right (Fig. 4g). This finding was further confirmed with ANOVA on LV1 scores which demonstrated that there were no cumulative significant differences between respective MMT conditions (Fig. 4h). Consideration of all these findings suggests encapsulated hMSCs yield a minimal therapeutic effect on subchondral bone in OA as this therapeutic yielded less bone thickening (volume) and hardening (attenuation). These disease modifying effects were similar to those found for articular cartilage analyses as further disease development from the time point of intervention was attenuated. Furthermore, encapsulated hMSCs again did not provide a restorative effect as there was still significant subchondral bone thickening (thickness) and bone hardening (medial 1/3 attenuation).

### Encapsulated hMSCs augmented osteophyte development in OA

One of the major associated phenotypes in OA is the development of tissues that form along the marginal edges of joints, known as osteophytes.^60^ These formations are the most common radiographic finding of OA in the clinic and therefore are a key consideration in studying therapeutics for this disease.^57,58,61^ Osteophytes consist of cartilaginous and mineralized portions, as they undergo an endochondral-like ossification process in formation, both of which were quantified in the current study in addition to a total osteophyte parameter (consisting of a combination of both cartilaginous and mineralized osteophytes).^60^

MicroCT analysis was implemented to quantitatively assess osteophyte volumes. Mineralized osteophyte volume was significantly greater in all MMT conditions relative to the Sham group (Fig. 5a). Furthermore, MMT/Encap hMSC and MMT/hMSC, which were not found to be different from one another, had significantly higher osteophyte volumes than the other two MMT conditions (Fig. 5a). Analysis of the other major osteophyte component (cartilaginous osteophytes) showed significantly higher volumes for all MMT groups relative to Sham (Fig. 5b). The MMT/Encap hMSC group also demonstrated increased cartilaginous osteophyte volumes relative to the MMT/hMSC and MMT/Empty Caps groups (Fig. 5b). Qualitative representation of these cartilaginous osteophytes can be viewed in the Saf-O histological images (Fig. 2f-j). Total osteophytes, a summation of mineralized and cartilaginous osteophytes, demonstrated similar findings to those for the individual parameters (Fig. 5c). A key difference identified was between MMT/Encap hMSC and MMT/hMSC, with the encapsulated condition demonstrating a significantly higher total osteophyte volume.

**Fig. 5.**
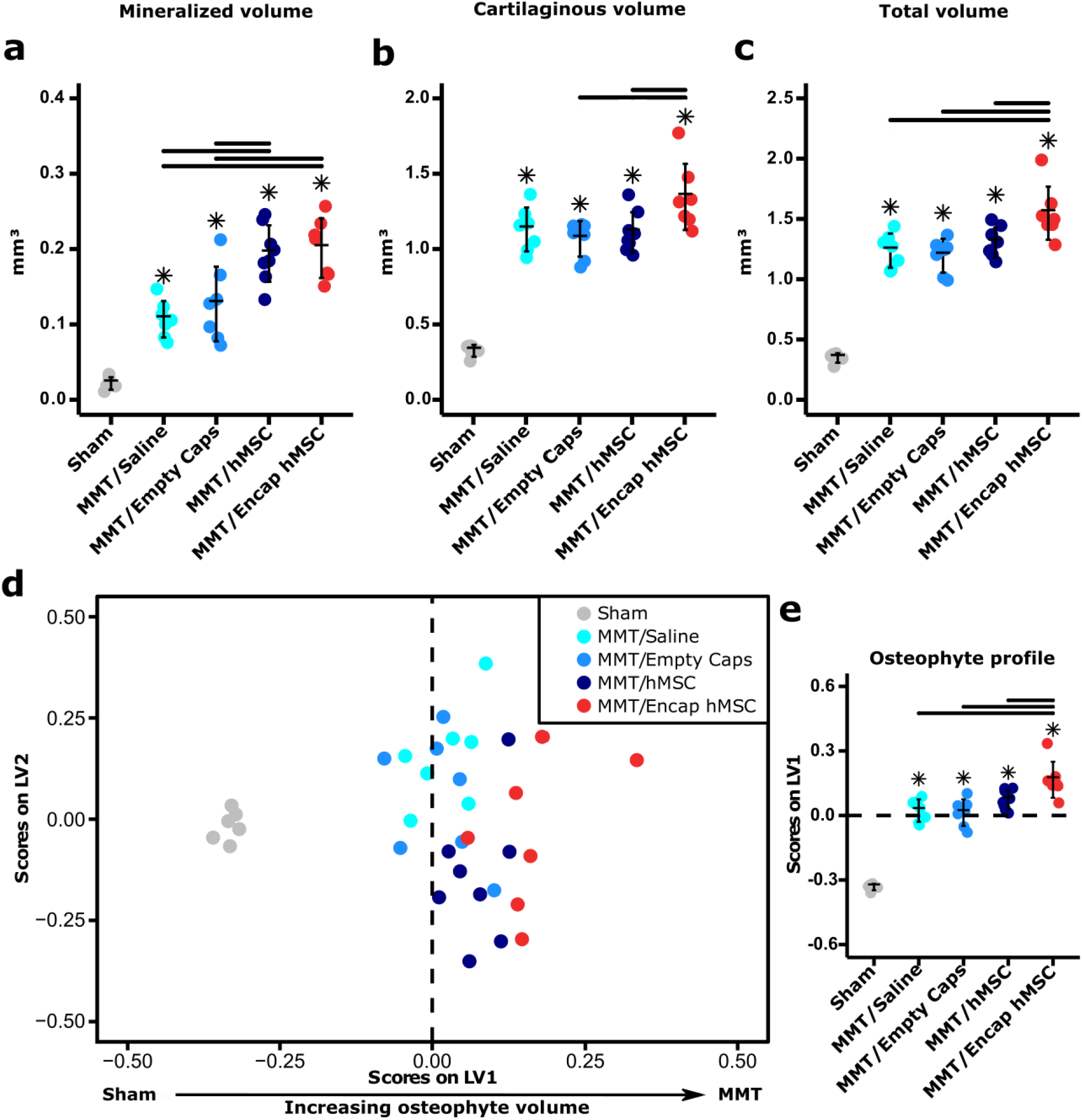
(a) Mineralized osteophyte volumes for all MMT groups were significantly greater than the Sham group; MMT/Encap hMSC and MMT/hMSC groups yielded significant increases in mineralized osteophyte volume compared to MMT/Saline and MMT/Empty Caps groups. (b) Cartilaginous osteophyte volumes for all MMT groups were significantly greater than the Sham group; MMT/Encap hMSC demonstrated significant increases in cartilaginous volume relative to MMT/hMSC and MMT/Empty Caps. (c) Total osteophyte volumes (Mineralized volume + Cartilaginous volume) for all MMT groups were again significantly greater than the Sham group; MMT/Encap hMSC yielded significantly greater total osteophyte volumes than all MMT conditions. (d) PLSDA analysis of overall osteophyte volumes depicted distinct separation of all groups based on osteophyte size, with Sham to the left and MMT/Encap hMSC to the right along LV1. (e) Statistical analysis of LV1 scores demonstrated significantly higher values for all MMT conditions, compared to Sham, with MMT/Encap hMSC showing increased volumes relative to all MMT conditions. Data presented as mean +/- SD. *n =* 6 for Sham, *n* = 7 for MMT/Saline, *n* = 7 for MMT/Empty Caps, *n* = 8 for MMT/hMSC and *n* = 7 for MMT/Encap hMSC. * represents significant differences (p < 0.05) between individual MMT conditions and Sham. Horizontal black bars indicate significance (p < 0.05) between individual MMT groups.

To assess the overall effect of encapsulated hMSCs on both cartilaginous and mineralized osteophytes, PLSDA established an LV1 that separated Sham to the left, and all MMT groups to the right based on increasing osteophyte volumes (Fig. 5d). ANOVA of LV1 scores displayed that all MMT conditions were significantly higher (increased volume) than Sham and that MMT/Encap hMSC was significantly higher than all other MMT conditions (Fig. 5e). Importantly, these results indicate that hMSCs, particularly the encapsulated hMSCs, potentiated osteophyte volumes relative to other MMT conditions that did not receive treatment. Even though increased osteophyte volumes have generally been viewed as an adverse outcome in OA, their development has been shown to occur independently of changes in articular cartilage morphology.^62^ Furthermore, osteophytes have been shown to increase motion segment resistance to both bending and compression forces, suggesting that osteophyte formation may reverse some of the mechanical stimuli that cause them to form, in a possible compensatory and protective role.^63^

### Biomaterial encapsulation of hMSCs induced a targeted paracrine response

The overall finding from the MMT study demonstrated a therapeutic effect of encapsulated hMSCs in preventing further cartilage degeneration in established OA. Another major finding drawn from the MMT study is the role that biomaterial encapsulation has in modulating the paracrine response of hMSCs *in vivo*. While the same cells (matched donor), administered at the same dose, were used for encapsulated and non-encapsulated therapeutic conditions, the two groups yielded differing levels of therapeutic efficacy. The encapsulated hMSCs elicited a more potent therapeutic effect compared to the non-encapsulated hMSCs which yielded a mild therapeutic effect in established OA. Specifically, non-encapsulated hMSCs demonstrated attenuation of increases in cartilage degeneration (volume and thickness) and subchondral bone remodeling (attenuation). These non-encapsulated hMSCs also yielded augmented osteophyte volumes, comparable to those yielded by encapsulated hMSCs. This led to the hypothesis that there would be significant differences in the secretome response to an OA microenvironment between non-encapsulated hMSCs and encapsulated hMSCs. Numerous studies have shown that biomaterials can alter MSC function, survival, and mechanotransduction; there is limited understanding of the effects encapsulation has on MSC cytokine expression which the current study sought to explore.^18,37,38,64^ To assess the effects of biomaterial encapsulation on the secreted cytokines from hMSCs in a simulated OA microenvironment, an *in vitro* cell culture model was used where the media was supplemented with the primary OA cytokine IL-1β. Cell viability immediately following encapsulation was 97.1 ± 3.1%, after which cells were plated. Cells were either conditioned in media alone (+CTRL) or stimulated with IL-1β in media (+IL-1β).^9,47,65^

Following treatment with or without IL-1β for 24 hours, media was collected and assessed for 41 immunomodulatory cytokines and chemokines (Fig. 6a). Background subtraction was performed for both non-stimulated and stimulated conditions; all cytokine values only show the cytokine levels resulting from hMSC paracrine expression. Both non-encapsulated and encapsulated hMSCs were responsive to IL-1β stimulation when compared to CTRL conditions. For non-encapsulated hMSCs, IL-1β stimulation yielded indiscriminate upregulation of all measured cytokines, compared to the non-encapsulated hMSC control (+CTRL). In contrast, IL-1β stimulation of encapsulated hMSCs yielded a more targeted response with distinct qualitative increases in six cytokines: IL-1β, IL-1RA, IL-7, IL-8, Granulocyte Colony Stimulating Factor (G-CSF), and IL-6, relative to the encapsulated hMSC control (+CTRL). PLSDA revealed LV1 and LV2 axes that separated differentially modulated cellular paracrine responses with encapsulated hMSC conditions separating on LV1 (+IL-1β separated left and +CTRL separated right) and non-encapsulated hMSCs separating along LV2 (+CTRL at the bottom and + IL-1β at the top of the axis; Fig. 6b). Significant separation of latent variable scores was confirmed with t-tests of both LVs, with Bonferroni correction applied, for encapsulated and non-encapsulated hMSCs on the LV1 and LV2 axes, respectively (Fig. 6c&d).

**Fig. 6.**
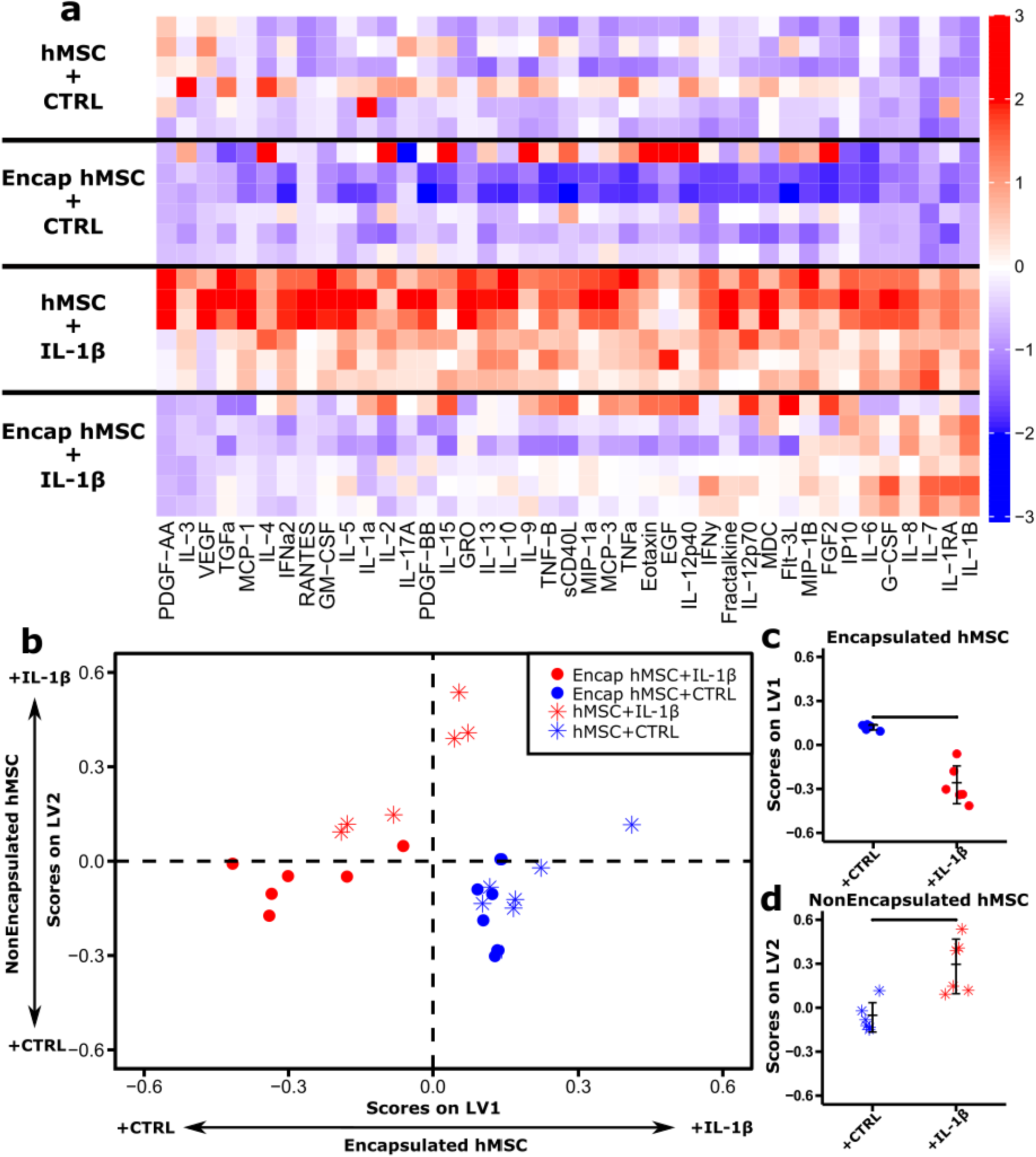
(a) Luminex analysis of 41 cytokines (columns; z-scored) secreted from hMSCs with (Encap hMSC) and without encapsulation (hMSC) in non-stimulated (+CTRL) and stimulated environments (+IL-1β; each row represents a single sample). (b) PLSDA analysis identified two profiles of cytokines, LV1 and LV2, that identified a distinct separation between treatment groups for both encapsulated and non-encapsulated hMSCs. (c&d) Independent analysis of scores on each of the respective latent variables demonstrated significant differences between non-stimulated (+CTRL) and stimulated environments (+IL-1β) for both encapsulated and nonencapsulated hMSCs. Data presented as mean +/- SD. *n =* 6 for all groups. Horizontal black bars indicate significant differences between non-stimulated (+CTRL) and stimulated (+IL-1β) groups.

To assess the effects of IL-1β stimulation on encapsulated cells, PLSDA was conducted on the encapsulated data alone (Fig. 7a). From LV1, the separation between CTRL to the left and IL-1β to the right can be easily observed. LV1 consisted of a profile of cytokines that correlated with the CTRL group (blue) or IL-1β treated cells (red; Fig. 7b). To determine which cytokines yielded differences in cytokine expression (Encap hMSC + CTRL vs. Encap hMSC + IL-1β), univariate analysis was performed on all cytokines that were upregulated with IL-1β stimulation (shown in red; Fig. 7b). Of the 18 cytokines that showed upregulation with IL-1β stimulation, eight were found to be significantly elevated, including the pro-inflammatory cytokines IL-1β, IL-6, IL-7 and IL-8, the anti-inflammatory cytokine IL-1RA and the chemokines G-CSF, macrophage derived chemokine (MDC; CCL-12), and interferon gamma-induced protein 10 (IP10; CXCL-10; Fig. 7c-j). These *in vitro* findings demonstrate a more modulated, and targeted response of encapsulated hMSCs when compared to the expression profile of non-encapsulated hMSCs, which yielded increased expression of all cytokines.

**Fig. 7.**
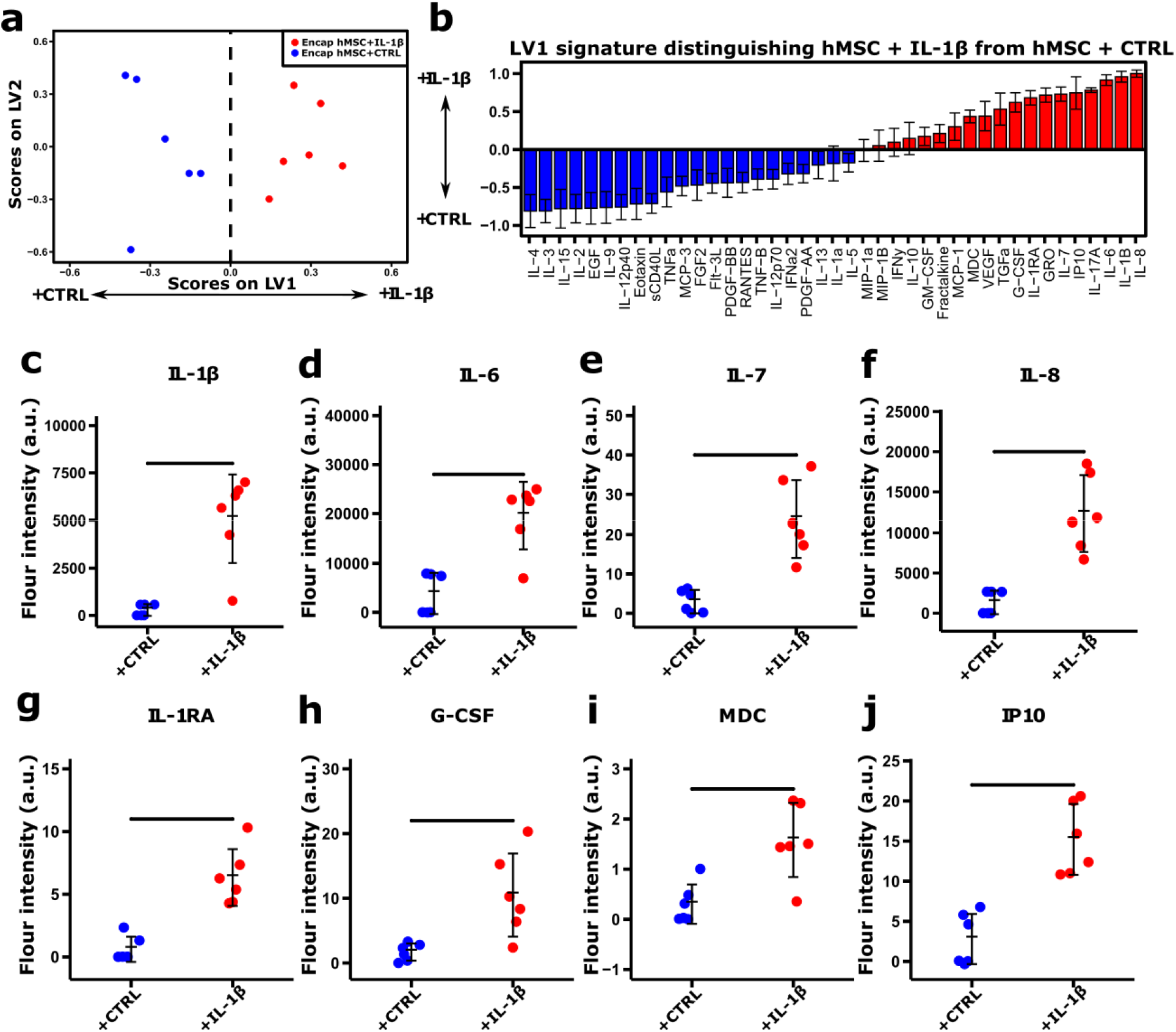
(a) PLSDA analysis of encapsulated hMSCs identified a single latent variable, LV1, that distinguished between Encap hMSC + CTRL on the left and Encap hMSC + IL-1β to the right. (b) The weighted profiles of cytokines showed relative expression of cytokines in CTRL conditions (blue) and IL-1β conditions (red). Error bars on each cytokine were computed by PLSDA model regeneration using iterative (1000 iterations) leave one out cross validation (LOOCV). (c-j) All measured cytokines that showed significant increased expression with IL-1β stimulation were assessed independently, using t-test with Bonferroni correction, for significance between CTRL and IL-1β conditions, with all significant findings presented. Encap hMSCs + IL-1β yielded increased expression in pro-inflammatory (IL-1β, IL-6, IL-7, IL-8), anti-inflammatory (IL-1RA), and chemotactic (G-CSF, MDC, IP10) cytokines. Data presented as mean +/- SD. *n* = 6 for all groups. Horizontal black bars indicate significant differences between non-stimulated (+CTRL) and stimulated (+IL-1β) groups.

The roles that cytokines play in OA pathology have been well documented, including a critical role for cytokines in osteophyte development.^13^ Previous studies have demonstrated that osteophyte growth is driven by cytokine release and not by mechanical forces on the joint capsule.^66–68^ However, a limitation of this study was that one of the major cytokines implicated in osteophyte development, TGF-β, was not part of the 41-plex cytokine panel. While previous studies have demonstrated that hMSCs yield an anti-inflammatory and regulatory paracrine expression profile in various disease states, the current study demonstrated increased pro-inflammatory cytokine secretion.^69,70^ Of note, IL-6 has been shown to decrease collagen II production and is implicated as a critical cytokine in subchondral bone remodeling in OA.^13,71^ Additionally, IL-7 has been demonstrated to increase production of MMP-13 and IL-8 has been shown to recruit additional neutrophils and further type II collagen degradation in OA.^72,73^ Even though these cytokines have been previously reported to be involved in OA pathogenesis when chronically elevated, it is important to note that encapsulated hMSCs yielded a more targeted response when compared to non-encapsulated hMSCs treated with IL-1β (Fig. 6a). Nonencapsulated hMSCs yielded increased expression of numerous other cytokines implicated in the OA inflammatory cascade (TNFα, IFNγ, IL-17).^71^ Furthermore, it is important to draw the distinction between the chronic nature of the OA inflammatory environment and the acute response induced by the hMSC secretome. To resolve chronic inflammation, an acute event is needed to bring in immune cells and activate different inflammatory cascades to resolve and induce a pro-regenerative response.^74^ This mechanism is common in other chronic inflammatory environments and wound healing environments where acute inflammatory events are necessary to resolve chronic inflammation and transition to pro-regenerative immune responses to regulate inflammation.^75–77^ Furthermore, the pro-inflammatory cytokines that showed increased expression in the current study (IL-6, IL-7, IL-8) have been implicated as significant mediators in wound healing and similar diseases that involve resolution of inflammatory events.^78–80^ Additionally, when looking at the duration of hMSC viability from our previous study, we observed that the hMSCs are only viable for the first ~9 days post-injection, thus the hMSCs likely respond to the local environment to help induce a local endogenous response which could have longer lasting therapeutic effects, particularly if it promoted the resolution of chronic inflammation.^18^

While encapsulated hMSCs elicited a pro-inflammatory response in the current study, they also secreted anti-inflammatory cytokines and chemokines which may have therapeutic potential. Specifically, the anti-inflammatory cytokine IL-1RA has been studied extensively in the context of OA with pre-clinical studies demonstrating a protective capacity on articular cartilage.^81,82^ Furthermore, a number of chemokines were increased (G-CSF, MDC, IP10) when stimulated with IL-1β, which would suggest that hMSCs could induce a response to recruit native stem and immune cells to the injury site. While the role of G-CSF has not been documented in OA, this cytokine is known to mobilize MSCs from bone marrow and has been found to promote cartilage repair in a pre-clinical full cartilage defect rabbit model.^83,84^ MDC (CCL-12), which is understudied in OA, has been shown to recruit memory T cells in patients with OA.^85^ The cytokine IP10 (CXCL-10) was also upregulated by encapsulated hMSCs. IP10 has been shown in previous studies to be specifically involved in recruitment of synovial macrophages in OA.^86^ While the cytokines of interest in the current study were categorized based on their most commonly identified pro-inflammatory, anti-inflammatory, or chemotactic nature it is important to note that cytokines are well known to have varying roles and can be both cell and context dependent.^87^ Most notably IL-6, which has both pro-inflammatory and antiinflammatory potential.^88,89^ While the present secretome assessment does not provide a direct link between specific cytokines and therapeutic efficacy *in vivo*, there were a number of potential cytokines that may be implicated in the therapeutic efficacy of hMSCs in OA. Further study of these specific cytokines, as well as their up-stream regulators and downstream targets, will provide new insights into the mechanisms involved in OA that may be therapeutically targeted with hMSCs. Additional investigation is merited to determine what characteristics of biomaterial encapsulation yielded this modulated response and whether further tailoring the biomaterial properties could further enhance the therapeutic efficacy of the hMSCs. This could include further study into the effects of the niche constructed by a 3D microporous hydrogel system (e.g. chemical composition, lack of peptide modification, pore size, topography) on hMSC paracrine function as all these properties have been demonstrated to directly impact the secretome of these cells.^90–92^

## Conclusions

In the current study, bone marrow derived hMSCs were encapsulated in sodium alginate microcapsules to study the effects of biomaterial encapsulation on the modulation of the paracrine signaling response and therapeutic efficacy of these cells in an OA microenvironment. The therapeutic potential of this cellular treatment was assessed in a pre-clinical rat model (MMT) of established OA, which is relevant because patients commonly seek treatment once OA is more readily evident and they have developed a more advanced stage of the disease. Encapsulation of hMSCs demonstrated a positive therapeutic effect by delaying further development of the disease; specifically, encapsulated hMSC treatment attenuated further cartilage degeneration and subchondral bone sclerosis. Though the encapsulated hMSCs provided a disease modifying protective effect, the treatment did not regenerate or restore the cartilage or subchondral bone back to levels comparable to Sham operated controls. These data suggest that the timing of hMSC treatment in the OA disease progression will be critical, as this treatment protected the integrity of the remaining tissue and thus suggests that treatment during earlier disease stages (when there is still tissue to protect) may have longer and more potent therapeutic effects. Though protective effects were observed on the cartilage and subchondral bone, encapsulated hMSCs yielded increased osteophyte volumes which have been identified as an unwanted phenotype for restoring joint function. The immunomodulatory potential of biomaterial encapsulation on hMSC function demonstrated a targeted paracrine response to a simulated OA microenvironment while non-encapsulated hMSCs showed an indiscriminate upregulation of all cytokines in the cytokine panel. While expression of numerous anti-inflammatory and regenerative cytokines were increased with hMSC encapsulation, there were also a number of pro-inflammatory cytokines that showed increased expression. In considering these latter findings, it is important to consider that this hMSC paracrine response is an acute response and that the secretion of these pro-inflammatory cytokines may be critical in resolving the chronic OA inflammatory environment. Together, the data from the current study demonstrated that biomaterial encapsulation of hMSCs modulated the paracrine response to a simulated OA microenvironment and enhanced the *in vivo* therapeutic efficacy of the hMSCs in preventing further disease progression in treating established OA.

## Acknowledgements

This work was supported in part by VA (SPiRE) Grant I21RX002372-01A1 from the United States (U.S.) Department of Veterans Affairs Rehabilitation Research and Development Service. The research was also supported in part by the DOD PRMRP Grant PR171379 and PHS Grant UL1TR000454 from the Clinical and Translational Science Award Program, National Institutes of Health, National Center for Advancing Translational Sciences. The authors would like to thank Mila Friedman for the histological analyses and Colleen Oliver for her support in the animal study.

## Disclaimer

The contents do not represent the views of the U.S. Department of Veterans Affairs or the United States Government.

