## Supplemental Figures for "Biomaterial encapsulation of human mesenchymal stromal cells modulates paracrine signaling response and enhances efficacy for treatment of established osteoarthritis"

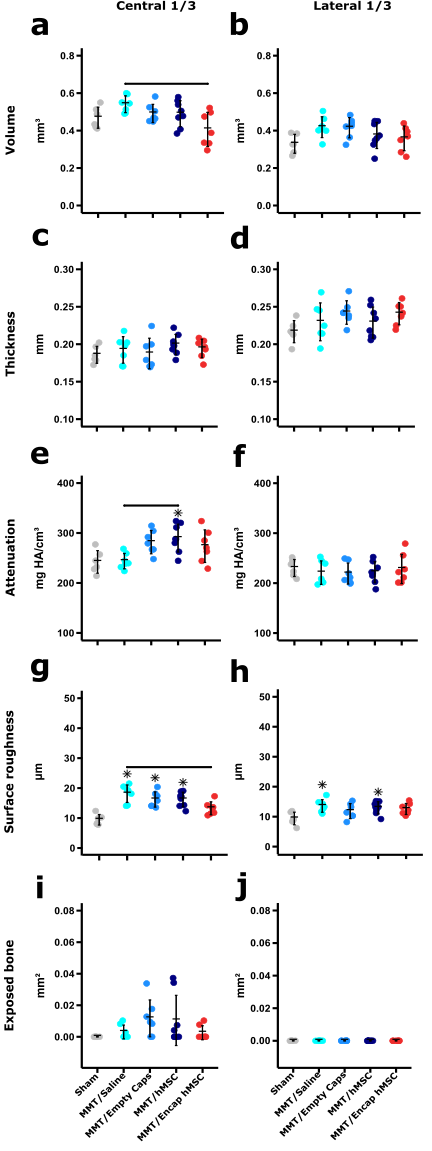

**Fig. S1.** (a) Central 1/3 articular cartilage volume yielded a single significant difference between MMT/Saline and MMT/Encap hMSC groups. (b) Lateral 1/3 articular cartilage volume yielded no significant differences between any groups. (c&d) Central 1/3 and lateral 1/3 articular cartilage thickness yielded no significant differences between groups. (e) Central 1/3 articular cartilage attenuation for the MMT/hMSC group was significantly greater than both the Sham and MMT/hMSC groups. (f) Lateral 1/3 cartilage attenuation showed no significant differences between any groups analyzed. (g) Lateral 1/3 cartilage surface roughness showed significantly higher values for the MMT/Saline, MMT/Empty Caps, and MMT/hMSC groups relative to Sham; the MMT/Encap hMSC group did show significantly less surface roughness than the MMT/Saline group. (h) Lateral 1/3 analysis of surface roughness yielded significantly increased surface roughness in the MMT/Saline and MMT/hMSC relative to Sham. (i) No significant differences were found between any groups for central 1/3 analysis of exposed bone surface area. (j) No groups showed any exposed bone in the lateral 1/3; no significant differences were found between any groups. Data presented as mean +/- SD. *n =* 6 for Sham, *n* = 7 for MMT/Saline, *n* = 7 for MMT/Empty Caps, *n* = 8 for MMT/hMSC and *n* = 7 for MMT/Encap hMSC. * represents significant differences (p < 0.05) between individual MMT conditions and Sham. Horizontal black bars indicate significance (p < 0.05) between individual MMT groups.

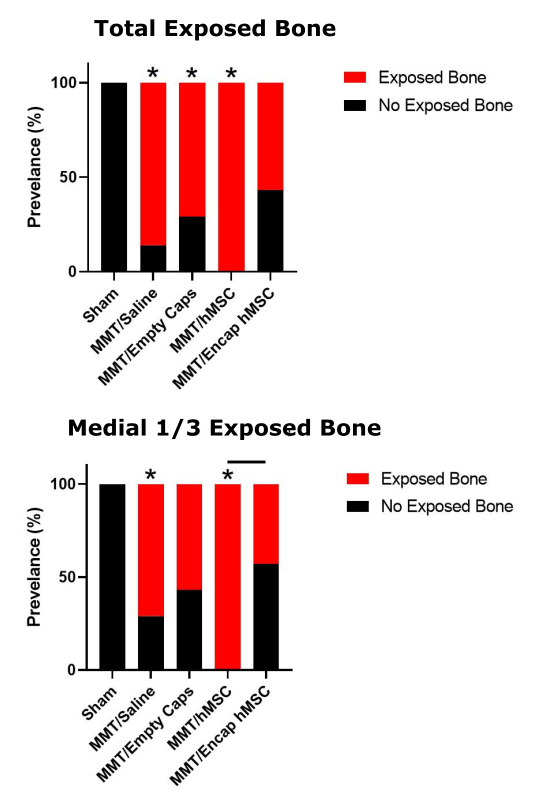

**Fig. S2.** (a) Incidence of exposed bone in the MMT/Encap hMSC group had the least number of samples with exposed bone (4/7) while all other MMT groups [MMT Saline (6/7), MMT/Empty Caps (5/7), and MMT/hMSC (8/8)] showed significantly more exposed bone incidence relative to Sham (0/6); no significant differences were found between any of the MMT groups. (b) Analysis of incidence of exposed bone on the medial aspect of the joint yielded increased exposed bone incidence in both the MMT/Saline (5/7) and MMT/hMSC (8/8) groups relative to Sham (0/6); no significant differences were found between Sham and MMT/Encap hMSC (3/7) or MMT/Empty Caps (4/7) groups. A single difference between MMT groups was found between the MMT/hMSC and MMT/Encap hMSC groups. Data presented as prevalence of exposed bone. *n =* 6 for Sham, *n* = 7 for MMT/Saline, *n* = 7 for MMT/Empty Caps, *n* = 8 for MMT/hMSC and *n* = 7 for MMT/Encap hMSC. * represents significant differences (p < 0.05) between individual MMT conditions and Sham. Horizontal black bars indicate significance (p < 0.05) between individual MMT groups.

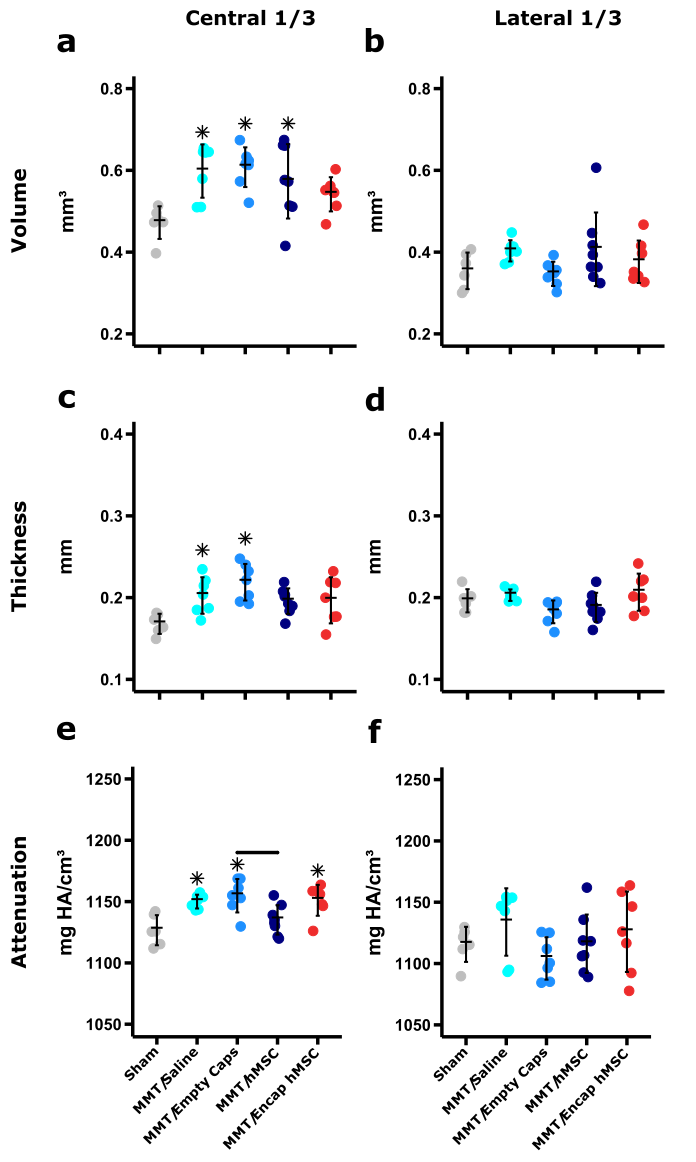

**Fig. S3.** (a) Central 1/3 subchondral bone volume for all MMT groups, except MMT/Encap hMSC, were significantly greater than the Sham group. (b) Lateral 1/3 subchondral bone volume yielded no significant differences between any groups. (c) Central 1/3 subchondral bone thickness analysis yielded significant increases in the MMT/Saline and MMT/Empty Caps groups relative to Sham; no significant differences were found between any of the MMT conditions. (d) No significant differences were found between any of the groups for subchondral bone thickness in the lateral 1/3 region. (e) Central 1/3 subchondral bone attenuation yielded significant increases for MMT/Saline, MMT/Empty Caps, and MMT/Encap hMSCs relative to Sham; a significant increase was also found between MMT/Empty Caps and MMT/hMSC. (f) In the lateral 1/3 region, no significant differences were found. Data presented as mean +/- SD. *n =* 6 for Sham, *n* = 7 for MMT/Saline, *n* = 7 for MMT/Empty Caps, *n* = 8 for MMT/hMSC and *n* = 7 for MMT/Encap hMSC. * represents significant differences (p < 0.05) between individual MMT conditions and Sham. Horizontal black bars indicate significance (p < 0.05) between individual MMT groups.
